# *In vivo* screening for toxicity-modulating drug interactions identifies antagonism that protects against ototoxicity in zebrafish

**DOI:** 10.1101/2023.11.08.566159

**Authors:** Ethan Bustad, Emma Mudrock, Elizabeth M. Nilles, Andrea McQuate, Monica Bergado, Alden Gu, Louie Galitan, Natalie Gleason, Henry C. Ou, David W. Raible, Rafael E. Hernandez, Shuyi Ma

## Abstract

Ototoxicity is a debilitating side effect of over 150 medications with diverse mechanisms of action, many of which could be taken concurrently to treat multiple conditions. Approaches for preclinical evaluation of drug interactions that might impact ototoxicity would facilitate design of safer multi-drug regimens and mitigate unsafe polypharmacy by flagging combinations that potentially cause adverse interactions for monitoring. They may also identify protective agents that antagonize ototoxic injury. To address this need, we have developed a novel workflow that we call Parallelized Evaluation of Protection and Injury for Toxicity Assessment (PEPITA), which empowers high-throughput, semi-automated quantification of ototoxicity and otoprotection in zebrafish larvae. By applying PEPITA to characterize ototoxic drug interaction outcomes, we have discovered antagonistic interactions between macrolide and aminoglycoside antibiotics that confer protection against aminoglycoside-induced damage to lateral line hair cells in zebrafish larvae. Co-administration of either azithromycin or erythromycin in zebrafish protected against damage from a broad panel of aminoglycosides, at least in part via inhibiting drug uptake into hair cells via a mechanism independent from hair cell mechanotransduction. Conversely, combining macrolides with aminoglycosides in bacterial inhibition assays does not show antagonism of antimicrobial efficacy. The proof-of-concept otoprotective antagonism suggests that combinatorial interventions can potentially be developed to protect against other forms of toxicity without hindering on-target drug efficacy.

## Introduction

Ototoxicity is a debilitating side effect of over 150 medications used to treat a broad range of conditions, including cancer and recalcitrant infections^1^. The sensorineural hearing or balance impairments that result from ototoxicity harm patient quality of life and incurs follow-up costs averaging $300,000-$1 million per patient^1^. Adverse drug-drug interactions (DDIs) with the potential to exacerbate ototoxicity complicate the implementation of treatment for multiple concurrent conditions. Currently, many DDIs are discovered only after the drugs have reached market^2^ as drug effects on the ear are not routinely tested in pre-clinical and clinical trials^3^. This late-stage discovery exacerbates the toll on patient health and financial costs. New approaches for preclinical identification of potential DDIs would mitigate unsafe polypharmacy by flagging regimens may increase toxicity for monitoring and even potentially identify protective agents that antagonize ototoxic injury.

Zebrafish (*Danio rerio*) are a well-established model organism for studying ototoxicity that offer the advantage of conserved vertebrate physiology, as well as compatibility with high-throughput assays typically limited to cell-based models^4–10^. Zebrafish share a high degree of genetic similarity with humans (70% of human genes have a clear zebrafish ortholog^11^), and the zebrafish lateral line hair cells (HCs), which detect vibrations in water, are structurally and functionally homologous to HCs of the human inner ear that sense vibrations of sound waves to enable hearing^12, 13^. Zebrafish larval lateral line HCs have functionality that more closely resembles that of mammalian HCs *in vivo* relative to cell lines derived from cochlear tissues, whilst also offering the potential for higher-throughput profiling experiments relative to *ex vivo* mammalian inner ear explant models that are laborious to establish^14^. Importantly, studies have shown that zebrafish larval lateral line HCs are sensitive to the same drugs that cause ototoxicity in humans^12, 13^, and that the majority of drug interactions tested in zebrafish have replicated in humans^7^.

The gold standard assay for ototoxic drug screening in zebrafish requires expert evaluation of multiple, individual neuromasts; attempts have been made to automate the analysis process^15, 16^, but throughput remains limited, preventing systematic evaluation of DDI. To address this challenge, we have developed a novel workflow that we call Parallelized Evaluation of Protection and Injury for Toxicity Assessment (PEPITA) for high-throughput, semi-automated quantification of ototoxicity and otoprotection in zebrafish larvae. By combining robotics-assisted 96-well plate-based microscopy imaging of whole zebrafish larvae and computational image analysis of the lateral line, our workflow empowers quantification of HC damage in hundreds of fish per day. Besides enabling larger-scale drug screening for putative ototoxic and otoprotective agents, PEPITA also enables the quantification of DDI outcomes of combinatorial drug co-administration.

By applying PEPITA to characterize ototoxic drug-drug interaction outcomes, we have discovered an antagonistic interaction between macrolide and aminoglycoside antibiotics that confers protection against aminoglycoside-induced damage to lateral line HCs in zebrafish larvae. Co-administration of either azithromycin or erythromycin in zebrafish protected against damage from a broad panel of aminoglycosides. Interestingly, co-administration of these macrolides with aminoglycosides in bacterial growth assays do not show a corresponding antagonism of antimicrobial efficacy. The proof-of-concept otoprotective antagonism between macrolides and aminoglycosides suggest that combinatorial interventions can be developed that protect against other forms of toxicity without hindering on-target efficacy. The platform PEPITA empowers the systematic screening for candidate combinations that elicit these protective interactions in the context of ototoxicity.

## Results

### High-throughput quantification of lateral line damage and ototoxic drug interactions with PEPITA

We have developed PEPITA, a novel, semi-automated workflow for streamlined quantification of ototoxicity in zebrafish (**Figure 1A, Figure S1**). The PEPITA workflow builds upon previous work that demonstrated that damage to zebrafish larval lateral line HCs could be quantified by evaluating the brightness intensity of fluorescently stained HCs using vital dyes such as YO-PRO-1 ^17–21^. Our platform enumerates relative residual brightness of YO-PRO-1-stained HCs post exposure of larvae to doses of individual or combinations of drugs. We quantify this remaining HC fluorescence as a proxy measure that is inversely proportional to the damage elicited by the ototoxic treatments.

**Figure 1.**
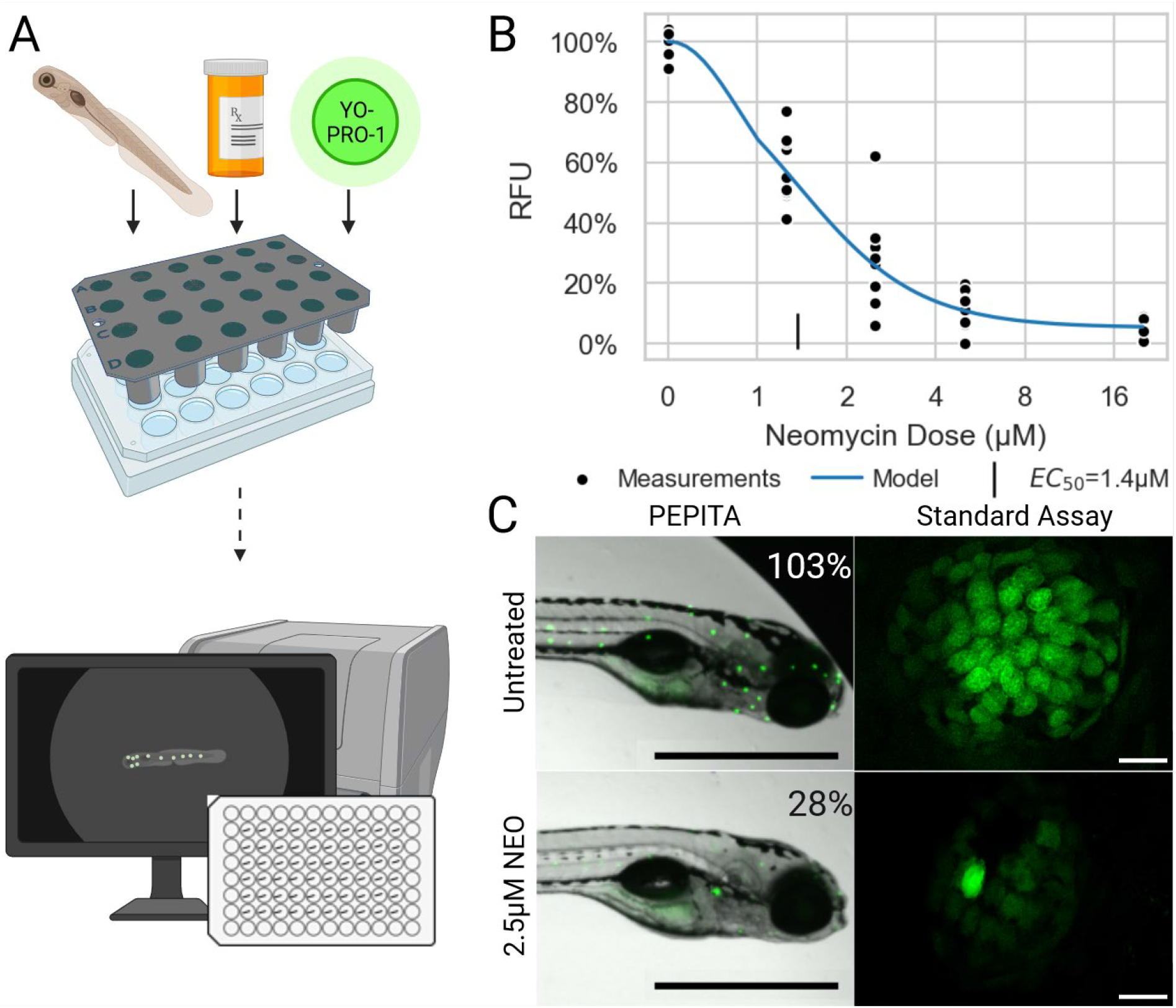
Overview of PEPITA workflow. (A) Zebrafish at 5dpf are placed into custom-made strainers in multi-well plates to be drug-treated and stained. After drug treatment for 4h and YO-PRO-1 staining for 20min, fish are anesthetized and imaged in brightfield and fluorescent channels. The resulting images are then quantified and analyzed. (B) A representative neomycin dose-response curve generated from the PEPITA workflow, with relative fluorescence units (RFU) at increasing neomycin concentrations. (C) A comparison between an image and quantification from PEPITA on the one hand, versus the gold standard high-magnification quantification assay on the other. Similar results are obtained via either method. Scale bars in whole-fish images represent 1 mm; in confocal, 10 µm. Figure created with BioRender.com

PEPITA enables quantification of hundreds of individual YO-PRO-1-stained fish per experiment. This high-throughput quantification of ototoxic damage in zebrafish larvae enables us to perform granular, reproducible characterizations of ototoxic dose-response curves for each drug of interest (**Figure 1B, Figure S2**).

To compare PEPITA’s ototoxic dose response characterization with the gold-standard approach of quantifying ototoxic damage ^12, 16, 17^, we also performed side-by-side characterizations with *myo6b::gfp* fish (**Figure 1C, Figure S2**). AB larvae stained with YO-PRO-1 and *myo6b::gfp* larvae present very similar neuromast appearance when untreated, at both high and low magnification, and decrease in fluorescence in a comparable way when treated with increasing doses of ototoxic drug. The two strains exhibit differences in doses necessary to achieve comparable levels of fluorescence inhibition. For instance, in our hands, the neomycin dose required to elicit 50% of maximal HC damage (EC_50_, see Methods for details) was 20μM in *myo6b::gfp* fish and 2.5μM in AB fish stained with YO-PRO-1 (**Figure S2, Figure S3**). Even in *myo6b::gfp* fish, this is lower than what has been observed previously, likely due to salinity differences, especially in calcium and magnesium, between our fish water and that of other investigators^22, 23^.

The shift between YO-PRO-1–stained AB and *myo6b::*gfp fish is likely due to a subtle difference in what is measured between the two: HC survival versus HC functionality. The *myo6b::gfp* fish express GFP in HC cytoplasm on the myosin 6 promoter, so stresses should not affect cell fluorescence up to the point of membrane rupture. In contrast, AB fish are stained with YO-PRO-1, which is taken up by the cell and fluoresces when it binds DNA ^24^. This means that, in addition to cell death, changes in uptake, trafficking, and DNA organization – all important signs of HC functionality – can affect fluorescence. Aminoglycosides cause damage at even relatively low doses, and at sufficient doses induce cell death by various pathways ^25^. While HC death would result in reduced fluorescent signal for both *myo6b::gfp* and AB fish, sub-lethal HC damage affecting cell functionality would only reduce fluorescence for YO-PRO-1-stained AB fish. This seems to explain the dose shift between the two strains. This implies an important caveat with regard to screening individual compounds with PEPITA: compounds that block YO-PRO-1 uptake without causing ototoxic damage will likely yield false positive indications for HC death when using AB fish stained with YO-PRO-1 alone, as we have observed with benzamil (data not shown). Complementary characterization with *myo6b::gfp* or other fish lines with transgenically labeled HCs will help to resolve these false positives. While our post-processing is currently optimized for reading out AB staining data, the platform is flexible to the use of other transgenic or stained fish as well.

Despite these differences in the underlying realities being measured, and the resulting difference in effective concentrations, observed interaction effects appear to be comparable. When detected, antagonism between drug pairs as quantified by windowed excess over Bliss (wEOB, see Methods for detailed description) is observed to comparable degrees in both assays (**Figure S4**).

### Azithromycin broadly antagonizes aminoglycoside ototoxicity and confers otoprotection

The high-throughput nature of PEPITA enables screening of ototoxic interactions experienced by fish by concomitant administration of combinations of drug treatments. In an initial pairwise screen of neomycin (NEO) against 13 compounds reported in the literature to impact HC survival and function ^1, 26–31^, we found that azithromycin (AZM) was one of the strongest antagonizing compounds to ototoxicity (**Figure S5**). To study the impact of this antagonism on the extent of otoprotection conferred by co-administration of AZM, we compared the extent of HC damage at varying doses of NEO exposure, with and without co-treatment with varying doses of AZM (**Figure 2A-B**). Although administration of high doses of AZM confers ototoxic damage (exposure to 380 µM AZM for 4 hours resulted in 48% residual HC fluorescence in YO-PRO-1 stained larvae, as quantified by relative fluorescence units (RFU), **Figure 2A**), co-administration of 96 µM of azithromycin rescued HC viability at concentrations of neomycin that were otherwise toxic to HC (exposure to 6.4 µM NEO for 4 hours resulted in 10% RFU in the absence of AZM, but 88% RFU when co-administered with 96 µM AZM, **Figure 2A**), and shifted the dose of neomycin needed to elicit 75% maximal HC damage (EC_75_) by 13-fold (3.3 µM to 42 µM, **Figure 2C**). This corresponded to wEOB score of −0.36, indicating strong antagonism (**Figure 2B**). This antagonism was observable in both AB and *myo6b::gfp* experiments (**Figure S4**).

**Figure 2.**
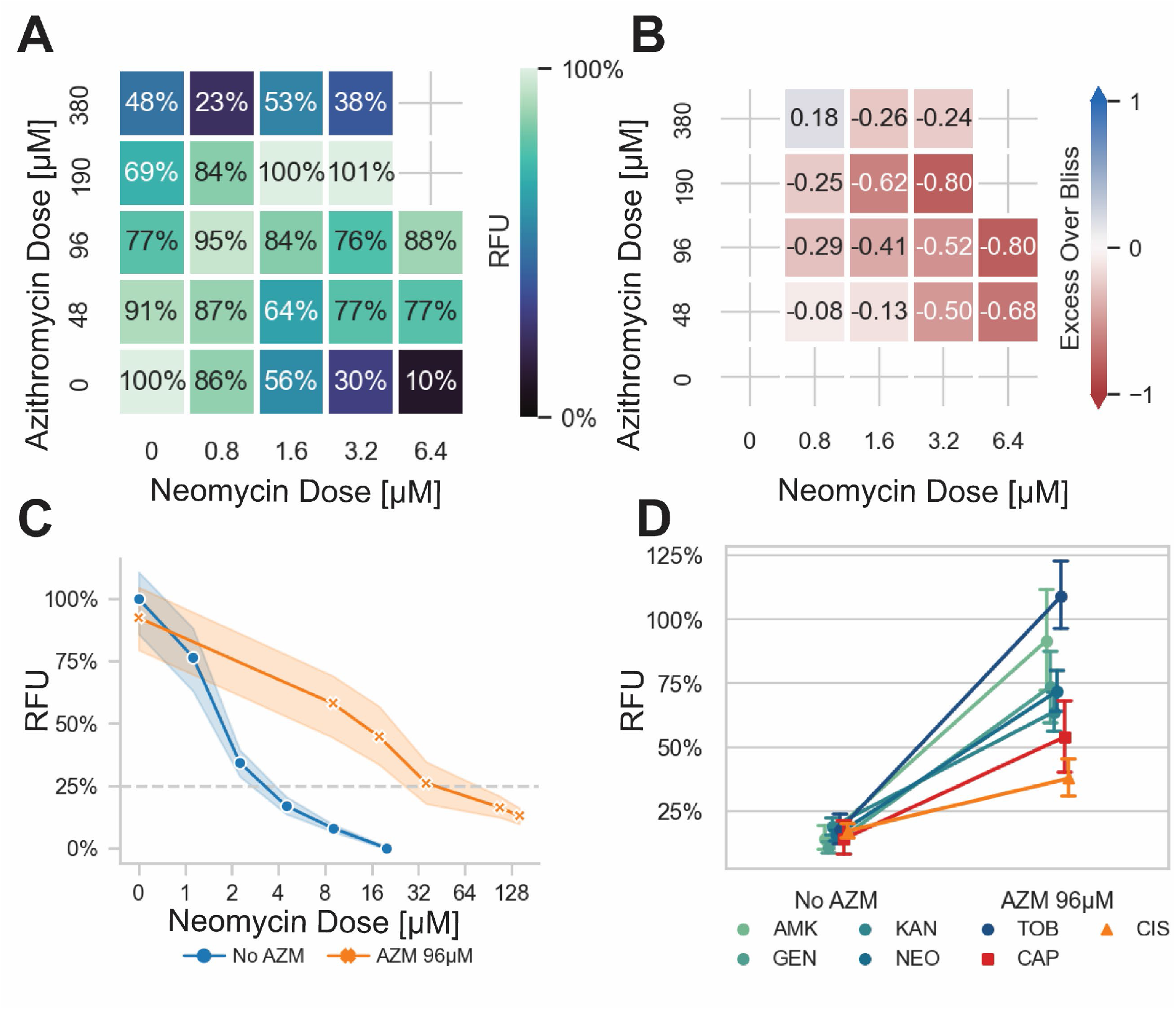
Azithromycin antagonizes aminoglycoside-induced ototoxicity in a zebrafish model. (A) Response seen when fish were exposed to increasing concentrations of azithromycin (AZM) and neomycin (NEO) in checkerboard format, as quantified by PEPITA in relative fluorescence units (RFU). Fish exposed to a combination of AZM and NEO experienced less ototoxic damage than those exposed to NEO alone, at all significantly ototoxic doses of NEO (i.e. ≥1.6μM) and AZM doses up through 190μM. In an extreme case, a dose of 6.4μM NEO, causing 90% HC damage (10% RFU), is reduced to minimal damage (88% RFU) by the addition of 96μM AZM. (B) excess over Bliss values calculated for the previous checkerboard data: positive numbers indicate synergy, negative numbers antagonism. The trend of reduced damage seen in the checkerboard translates to consistent antagonism, with an overall wEOB of −0.36 for this experiment. (C) Comparison of lateral line HC ototoxic dose response elicited by NEO with vs. without AZM co-administration (96 μM). (D) Comparison of lateral line HC damage in response to ototoxic drug exposure with vs. without AZM co-administration (96 μM). Drugs tested: AMK = amikacin, GEN = gentamicin, KAN = kanamycin, NEO = neomycin, TOB = tobramycin, CAP = capreomycin, CIS = cisplatin. Doses of drugs tested were selected to elicit 70% - 95% hair cell damage in the absence of AZM co-administration. Aminoglycosides as a group are more antagonized than the non-aminoglycoside drugs tested (p < 0.0001).

We wondered at the extent to which the antagonistic interaction between AZM and NEO observed with ototoxic response generalized to other aminoglycosides. We therefore tested ototoxic interactions between AZM and 5 aminoglycosides (neomycin (NEO), amikacin (AMK), gentamicin (GEN), tobramycin (TOB), and kanamycin (KAN)) (**Figure 2D, Figure S6**). Among interactions with AZM, co-administration with each of the 5 aminoglycosides exhibited otoprotective antagonistic interactions. For doses of each aminoglycoside that reduced functional HC survival by at least 70% when treated alone, co-administration with AZM resulted in 62%pt. protection in HC survival on average (improving from 15% survival to 77% survival on average; **Figure 2D**). In comparing the ototoxic HC damage dose response, the EC_75_ of each aminoglycoside shifted at least 8-fold with AZM co-administration (**Table 1, Figure S7**).

**Table 1.** Magnitude of EC75 shift when ototoxic drugs are co-administered with a macrolide as compared to administered alone. The aminoglycosides display large shifts with high levels of statistical significance; CAP and CIS show lower shifts that border on significance. The *p*-values listed represent the probability that the dose-response curve shift we observed occurred solely due to chance, as quantified by an extra sum-of-squares F test.

| Macrolide | Ototoxin | EC75 Shift | Shift *p*-value |
| --- | --- | --- | --- |
| Azithromycin (AZM) | Amikacin (AMK) | 8-fold | 0.0003 |
|  | Gentamicin (GEN) | 13-fold | <0.0001 |
|  | Kanamycin (KAN) | >16-fold | <0.0001 |
|  | Neomycin (NEO) | 13-fold | <0.0001 |
|  | Tobramycin (TOB) | >16-fold | <0.0001 |
|  | Capreomycin (CAP) | 2.8-fold | 0.057 |
|  | Cisplatin (CIS) | 1.8-fold | 0.048 |
| Erythromycin (ERY) | Neomycin (NEO) | 8-fold | <0.0001 |

We also tested the impact of AZM co-administration on protection of ototoxicity with neomycin (NEO) and gentamicin (GEN) at two different time points: 1 hour (acute) and 4 hours (extended) of treatment. These timepoints were selected because they had previously been shown to convey differences in protection between NEO and GEN when co-administered with different cell death inhibitor compounds ^25^. We found that AZM elicited significant protection against both NEO and GEN after 1 hour and 4 hours of treatment (for all conditions in which monotherapy induced a significant level of damage, AZM co-treatment conferred a significant level of otoprotection, **Figure S8A-D**).

### Azithromycin-induced otoprotective antagonism is significantly pronounced with aminoglycosides

To evaluate the specificity of AZM-mediated otoprotective antagonism, we measured the ototoxic interactions of AZM coadministration with non-aminoglycoside drugs that convey ototoxicity clinically and share similarities in their modes of toxicity and action: cisplatin and capreomycin. Cisplatin (CIS) was selected because of the similarity of the drug uptake mechanism in HCs relative to aminoglycosides (both sets of drugs require mechanoelectrical transduction (MET) channel activity for HC uptake) ^14^. Capreomycin (CAP) was selected because it is an ototoxic antibiotic with a similar mode of action compared to aminoglycosides, but has a chemical structure with distinct chemical properties ^32–34^.

We found that AZM co-administration conferred significant but less pronounced protection from CAP-associated damage, (**Figure 2D, Figure S7**). Exposure to 25μM CAP yielded 90% HC inhibition in the absence of azithromycin (CI 83%-96%) and 60% HC inhibition in the presence of azithromycin co-administration (CI 45%-76%) (**Figure 2D**). In comparing the ototoxic HC damage dose response, the EC_75_ of CAP shifted 2.8-fold (**Table 1, Figure S7**). Notably, this protection only seems to be present in highly damaging doses of CAP as the EC_50_ of CAP shifted 1.9-fold (CI −1.5 to 3.8-fold).

Similarly, AZM co-administration conferred significant but modest protection against damage caused by higher doses of CIS. Exposure to 600μM CIS yielded 81% HC inhibition (CI 78%-83%) in the absence of AZM and 57% HC inhibition (CI 46%-68%) in the presence of AZM co-administration. As with CAP, AZM co-administration conferred stronger protection under conditions of high HC damage (EC_75_ of CIS shifted 1.8-fold (CI 1.4 to 2.2-fold)) than lower HC damage (EC_50_ of CIS shifted 1.4-fold, CI −1.9 to 2.8-fold) (**Table 1, Figure S7**).

### Macrolide antibiotics broadly confer antagonistic otoprotection against aminoglycoside ototoxicity

To evaluate the extent to which AZM’s otoprotective antagonism against aminoglycosides extends to other drugs in its class, we also tested two other commonly used macrolide antibiotics, erythromycin (ERY) and clarithromycin (CLM), on drug interactions with aminoglycosides. Based on our preliminary experiments, we found antagonistic interactions between CLM and AMK, GEN, KAN, and NEO (**Figure S6**); however, we estimated that the optimally protective dose of CLM would be higher than the solubility threshold in water, so we chose to focus our characterization on ERY. We found that ERY also conveyed antagonistic otoprotection broadly against aminoglycoside ototoxicity during co-administration, although the most otoprotective dose of ERY was significantly higher than AZM (**Figure S7**). The EC_75_ of NEO shifted 8-fold (CI 4.0 to 17-fold) with co-administration of ERY (**Table 1, Figure S7**).

### Macrolides interfere with aminoglycoside uptake into hair cells

Given that macrolide co-administration appeared to confer comparable otoprotection against neomycin after both acute (1-hour) and extended (4-hour) exposure, we hypothesized that the mechanism of macrolide-associated protection involved influencing drug uptake.

To test the effect of macrolide co-administration on aminoglycoside uptake, we adopted two complementary approaches: (1) measuring the effect of AZM administration on MET channel activity via HC uptake of FM1-43, a dye whose transport is mediated by MET channel^35^ (**Figure 3A**); and (2) testing the effect of AZM co-administration on HC uptake of fluorescently-conjugated neomycin (**Figure 3B**).

**Figure 3.**
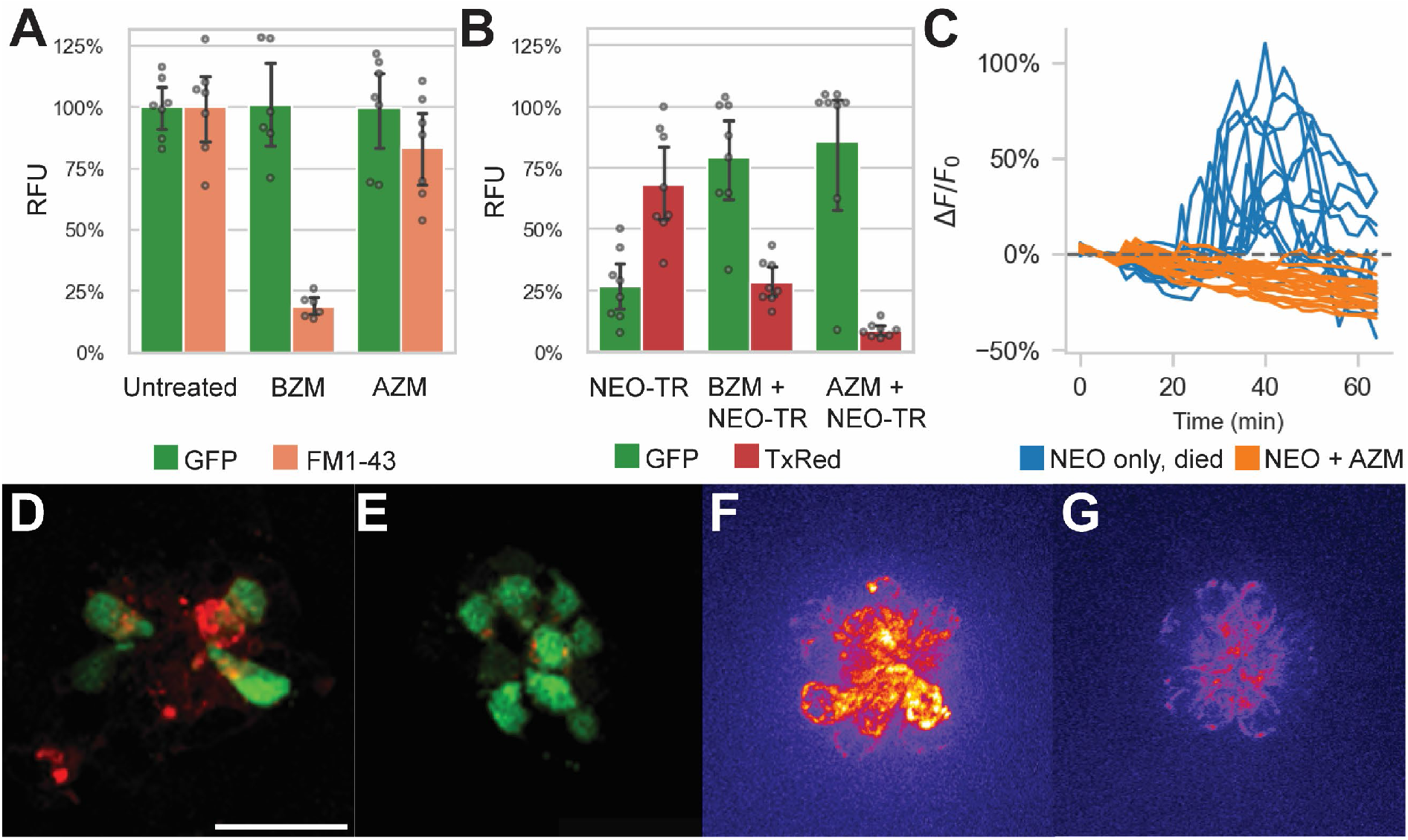
Impact of azithromycin and neomycin treatment on hair cell function. (A) AZM (190 µM) does not significantly hinder FM1-43 uptake into lateral line HCs (p = 0.15), suggesting it does not inhibit MET function. In contrast, administration of benzamil (BZM; 50 µM) as a positive control does significantly hinder FM1-43 uptake (p < 0.0001). (B) Effect of AZM (190 µM) co-administration on accumulation of TexasRed-conjugated neomycin (NEO-TR, 50μM exposure) within myo6b::gfp lateral line HCs. BZM (50 µM) co-administration with NEO was also evaluated as a positive control, and both conferred inhibition of NEO-TR uptake into the hair cells (BZM, p = 0.001; AZM, p = 0.0001). (D, E) Fluorescence microscopy images of the colocalization of NEO-TR (red) with GFP-expressing HCs (green), treated with NEO-TR alone (D) or with both NEO-TR and AZM (E). (C) Quantification of mitochondrial Ca^2+^ levels in individual HCs in response to NEO treatment alone (blue) or NEO plus AZM co-administration (orange), as measured by mitoGCaMP fluorescence signal. Drugs were administered at t = 10 minutes. (F, G) Fluorescence microscopy images of mitoGCaMP3 neuromasts at 30 minutes post drug administration, treated with NEO alone (F) or with both NEO and AZM (G). Scale bar for panels D-G represents 20µm.

We measured accumulation of FM1-43 in lateral line HCs in the presence vs. absence of AZM (190 µM) in *myo6b::gfp* larvae. AZM administration for 30 minutes prior to and during FM1-43 administration did not significantly change FM1-43 accumulation in HCs, whereas administration with the positive control, benzamil (BZM, 50 µM, a known MET channel blocker ^31^), for the same period showed significant inhibition of FM1-43 uptake (**Figure 3A**). These data suggest that MET channel activity is not impaired by AZM administration.

We also measured accumulation of TexasRed-conjugated neomycin (NEO-TR, 50 µM) in lateral line HCs in the presence vs. absence of AZM (190 µM) in *myo6b::gfp* larvae. AZM co-administration yielded reduced HC accumulation of NEO-TR (indicated by reduced TexasRed labeling in HCs after 30 minutes of exposure to TexasRed-conjugated drug), which was concomitant with reduced HC damage (indicated by increased GFP labeling in the same HCs) (**Figure 3B, 3D-E**). These data suggest that neomycin uptake is inhibited by azithromycin co-administration.

Should macrolides block aminoglycoside uptake into HCs, we expected downstream cell signaling pathways associated with aminoglycoside exposure would also be blocked. For example, it has been previously shown that once taken up by HCs, NEO induces a large spike in mitochondrial calcium, and subsequently in cytoplasmic calcium in HCs that go on to die ^36^. To test if AZM blocks these effects, we measured HC mitochondrial and cytoplasmic calcium responses in response to both NEO alone (50 µM) and co-administration of NEO (50 µM) with AZM (190 µM) in *mitoGCaMP3; cytoRGECO* larvae. Consistent with the previous findings, we demonstrated NEO induced an increase in mitochondrial (**Figure 3C, 3F, Figure S9**; n = 4 fish, 9 neuromasts) and cytoplasmic (not shown) calcium in HCs that went on to die (66% of those analyzed). There were no such increases in cells that survived. Co-administration of NEO with AZM, however, blocked all increases in mitochondrial calcium (**Figure 3C, 3G, Figure S9**; n = 5 fish, 10 neuromasts), with no changes in cytoplasmic calcium. Although we occasionally observed increases in mitochondrial calcium and low levels of cell death with AZM administration alone (**Figure S9**), there was no cell-death observed with NEO and AZM co-administration. These results suggest that AZM co-administration and subsequent NEO uptake inhibition further blocks the downstream signaling pathways associated with NEO-induced death, namely changes in HC calcium handling.

### Macrolides do not antagonize antimicrobial activity of aminoglycosides

Given that both the macrolides and aminoglycosides that we tested are antibiotic agents, we investigated whether they antagonized each other’s antimicrobial activity during co-administration. Checkerboard assay experiments in *Staphylococcus aureus, Escherichia coli* and *Mycobacterium abscessus* (representing gram-positive, gram-negative, and atypical bacteria, respectively), between AZM and 5 separate aminoglycosides (amikacin (AMK), gentamicin (GEN), kanamycin (KAN), neomycin (NEO), and tobramycin (TOB)) showed no pattern of antagonism (**Figure 4**). Average windowed excess over Bliss scores were 0.02 in *E coli*, 0.03 in *M. abscessus*, and 0.07 in *S. aureus*, suggesting on average little interaction to even slight synergy. In comparing the interaction of these aminoglycosides with AZM, for AMK this represents a 40%pt. shift toward antagonism in lateral line HC damage as compared to inhibition of these bacterial species on average; for GEN 41%pt. toward antagonism, for KAN 37%pt. toward antagonism, for NEO 42%pt. toward antagonism, and for TOB 27%pt. toward antagonism. These values suggest that treating a bacterial infection with a regimen comprising AZM and any of these aminoglycosides would cause on average less ototoxicity relative to treatment with the aminoglycoside alone, without antagonizing the antimicrobial activity of the antibiotics.

**Figure 4.**
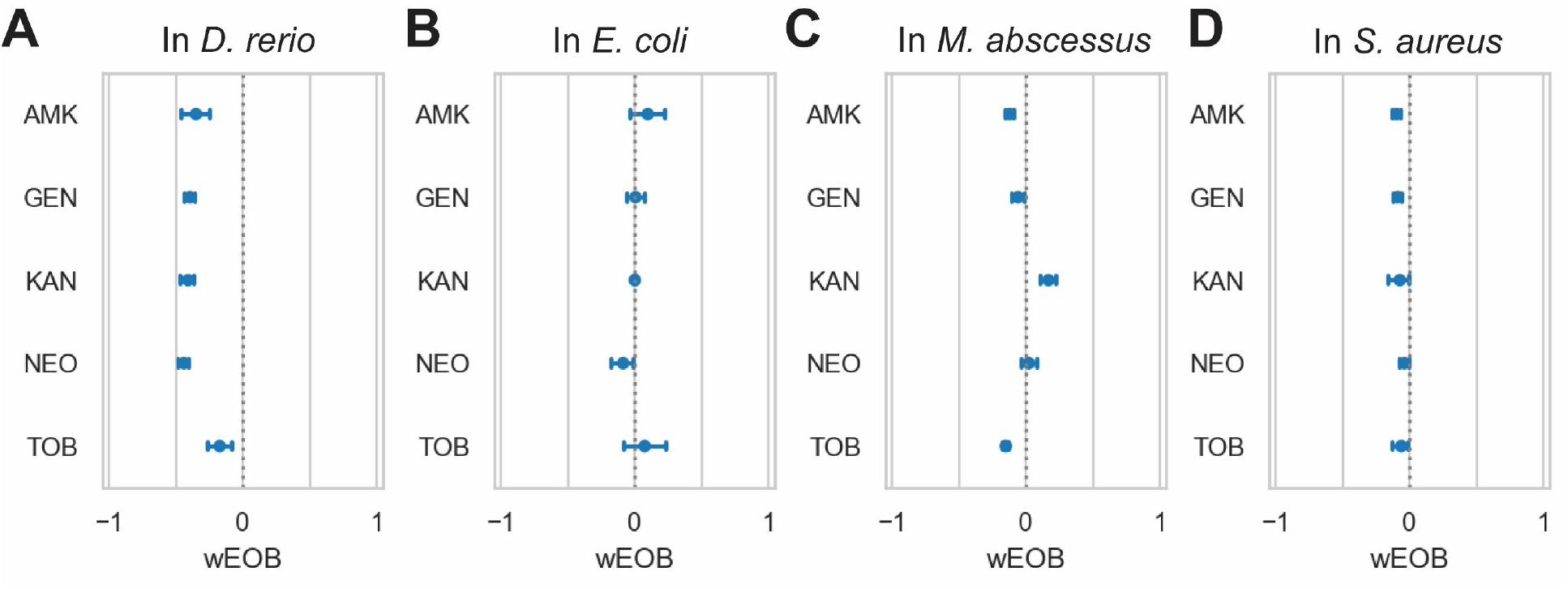
Azithromycin and aminoglycosides do not interact antagonistically in bacteria. (A) depicts ototoxic interactions in zebrafish as measured by PEPITA, while the subsequent three panels depict interactions as measured by checkerboard assay with the given bacterium, respectively: (B) E. coli, (C) M. abscessus, and (D) S. aureus. The x-axis quantifies the windowed excess over Bliss metric (wEOB), which measures drug interactions with negative numbers indicating antagonism and positive numbers indicating synergy. The y-axis contains 5 aminoglycoside antibiotics. None of the measured combinations yield a consistently antagonistic interaction in terms of the bacterial growth inhibition achieved, in contrast with the significant antagonistic protection AZM confers against damage induced by all of these drugs.

## Discussion

Given the multiplicity of potential drug-drug interactions and the complexity of their molecular consequences, anticipating the impacts of these interactions on toxicity outcomes is currently an important unresolved gap in the drug development process. High-throughput quantification of toxicity outcomes *in vivo* by tools such as PEPITA enables a unique lens into the treatment impact of combinations with toxicity-protective or toxicity-potentiating properties. By identifying candidate otoprotective multi-drug interventions, such as co-administration of macrolides with aminoglycosides, PEPITA can prioritize these combinations for follow-up translational preclinical studies. Since PEPITA can also quantify multiple drug treatment conditions in a single ototoxicity profiling experiment, PEPITA can also streamline investigation into the molecular mechanisms underlying toxicity and protection.

Previously published advances in automated quantification of ototoxicity in zebrafish have successfully streamlined the assessment of damage lateral line HCs from high-resolution images of YO-PRO-1-stained neuromast HCs taken by confocal microscopy ^16^. While this automated analysis approach facilitated faster interpretation of images once collected, the experimental workflow for preparing the treated and stained fish for imaging remained a laborious and time-consuming multi-day process, which also restricted the number of neuromasts captured to a relatively small number of representative neuromasts (7 per fish) on a relatively small number of fish. In contrast, the PEPITA workflow enables semi-automated image capture and analysis of live zebrafish larvae in a 96-well plate setting, without fixation or mounting of the larvae. By aggregating the staining information across multiple neuromasts from a 2x magnification image of each whole fish, the time required to acquire the necessary images for quantification is drastically reduced, typically lasting approximately 30 minutes per plate of 60 fish. Though some of the morphological details of the HCs are lost with a low-magnification image capture, PEPITA can capture more neuromasts per fish and more fish per experiment, which empowers us to achieve greater statistical power in assessing differences in ototoxic outcomes for each treatment.

Using PEPITA to characterize ototoxic drug-drug interaction outcomes, we have discovered a tissue-specific antagonistic interaction between macrolide antibiotics and aminoglycoside antibiotics that confers protection against aminoglycoside-induced injury to lateral line HCs in zebrafish larvae. Co-administration of macrolide antibiotics including azithromycin and erythromycin protected against lateral line HC damage induced by a panel of aminoglycosides. The successful identification of these protective interactions highlights the potential clinical translational utility of *in vivo* screening multidrug combinations for putative toxicity-protective interventions. Antagonism represents an understudied and underexploited modality of toxicity protection that might enable a novel avenue of repurposing clinically approved drugs towards toxicity-protective applications. Moreover, studying the molecular mechanisms underlying the protection of these antagonistic interactions might uncover new intervention targets that could protect against toxicity.

While clinical reports have noted observations of possible ototoxicity, the effect of macrolide antibiotics on HCs, including azithromycin, erythromycin, and clarithromycin, remain largely uncharacterized in the zebrafish lateral line model ^34^. Previous investigations with the mTOR inhibitor macrolide, rapamycin, have found that rapamycin confers significant protection against cisplatin ototoxicity by activating autophagy ^37^. Macrolide antibiotics including azithromycin are known to inhibit autophagy ^38, 39^, and the protection they confer against cisplatin ototoxicity is modest. This therefore suggests that their mechanism of protection is distinct from rapamycin. Although we observed evidence of ototoxic HC inhibition with high-dose azithromycin alone, our experiments with *myo6b::gfp* and *mitoGCaMP3* fish revealed that lower concentrations of macrolides inhibited aminoglycoside-induced HC toxicity when co-administered. This protective effect of macrolide antibiotics was similar when co-administered with aminoglycosides previously shown to have differences in their ototoxic mechanisms, such as gentamicin and neomycin ^25^, though the protective effects of macrolides on other ototoxic drugs that share some commonalities in mechanism, including cisplatin and capreomycin, are different and substantially reduced.

In the case of neomycin, the otoprotection conferred by macrolide co-administration is correlated with reduction in aminoglycoside uptake by HCs but not with impaired MET channel activity. MET channel activity has previously been shown to be required for aminoglycoside ototoxicity ^40, 41^. Our data suggest that there exist MET channel-independent modalities of impairing neomycin uptake in HCs and subsequent protection. Our findings are also consistent with an aminoglycoside uptake model in which aminoglycoside uptake enters the stereocilia of HCs through a mechanism other than direct MET channel transport, but relies on MET channel activity for transport into the HC body. This model is consistent with studies that have implicated members of the transient receptor potential cation channels (TRPs) in aminoglycoside uptake ^42–44^; others have also speculated that non-selective cation channels, such as connexins, pannexins, and P2X channels may also be involved in aminoglycoside uptake ^45^. Interestingly, the otoprotective macrolide antibiotics that we have identified in our study are also distinct in their chemical structures from known MET channel inhibitors that confer otoprotection (**Table S1**); this suggests that the aminoglycoside uptake inhibition and subsequent protection conferred by macrolides represents a novel biochemical strategy of otoprotection.

In comparing the drug interaction outcomes measured from co-administration of macrolides and aminoglycosides in zebrafish lateral line HCs versus interactions from bacterial inhibition assays with multiple species, we find that the interactions of these drugs are not broadly correlated across tissues and across the drug classes. Notably, most of the drug combinations did not exhibit antagonism when inhibiting *E. coli, S. aureus, or M. abscessus*. In contextualizing this antimicrobial efficacy along with the pronounced protection of HCs, our data suggest that macrolide co-administration might shift the therapeutic index of aminoglycosides at least 8-fold, when considering zebrafish HC ototoxicity. The tissue specificity of drug interaction outcomes is consistent with previous reports that drug-drug interaction outcomes are a condition-specific phenotype ^46^, and it also suggests that it may be possible to design regimens that improve therapeutic indexes by tuning antimicrobial interaction outcomes independently from host-targeting toxicity interaction outcomes. Given that differences exist between drug uptake and damage mechanisms in zebrafish lateral line HCs and adult mammalian inner ear HCs ^47^, additional studies are warranted to evaluate the extent to which otoprotective antagonism between macrolides and aminoglycosides also occurs in mammals.

The proof-of-concept otoprotective antagonism discovered by the PEPITA platform raises hopes for identifying analogously protective combinatorial interventions that do not hinder on-target efficacy for other toxic drugs. Although PEPITA is currently optimized to quantify ototoxicity, the underlying platform and image analysis pipeline could be adapted to investigate interactions with other organ toxicities assayable in the zebrafish larval system.

## Materials and methods

### Key resources table

| Reagent type (species) or resource | Designation | Source or reference | Identifiers |
| --- | --- | --- | --- |
| Strain, zebrafish (*Danio rerio*) | AB | ZIRC | http://zebrafish.org/fish/lineAll.php?OID=ZDB-GENO-960809-7 |
| Strain, bacteria | Escherichia coli ATCC 25922 | ATCC | https://www.atcc.org/products/25922 |
| Strain, bacteria | Mycobacterium abscessus ATCC 19977 | ATCC | https://www.atcc.org/products/19977 |
| Strain, bacteria | Staphylococcus aureus ATCC 29213 | ATCC | https://www.atcc.org/products/29213 |
| Genetic reagent (*Danio rerio*) | Tg(*myo6b:EGFP*)<sup>w119Tg</sup>; *myo6b::gfp* | 48 | https://zfin.org/ZDB-ALT-170321-13 |
| Genetic reagent (*Danio rerio*) | Tg(*myo6b:mitoGCaMP3*)<sup>w119Tg</sup>; *mitoGCaMP3*; | 49 | https://zfin.org/ZDB-ALT-141008-1 |
| Genetic reagent (*Danio rerio*) | Tg(*myo6b:RGECO1*); *cytoRGECO* | 50 | https://zfin.org/ZDB-ALT-150114-2 |
| Chemical compound, drug | Amikacin sulfate | Sigma-Aldrich | PHR1860-1G |
| Chemical compound, drug | Aspirin | ThermoFisher | 18-600-802 |
| Chemical compound, drug | Azithromycin dihydrate | ThermoFisher | AAJ6674006 |
| Chemical compound, drug | Benzamil | Sigma-Aldrich | B2417-50MG |
| Chemical compound, drug | Capreomycin sulfate | Sigma-Aldrich | PHR1716-500MG |
| Chemical compound, drug | Cisplatin | Biotechne | 2251 |
| Chemical compound, drug | Clarithromycin | TCI | C2220 |
| Chemical compound, drug | Cyclodextrin | ThermoFisher | AAH3113306 |
| Chemical compound, drug | Cyclosporine A | ThermoFisher | AAJ6319106 |
| Chemical compound, drug | Dexamethasone | ThermoFisher | AAA1759003 |
| Chemical compound, drug | Dihydrostreptomycin | ThermoFisher | ICN19452805 |
| Chemical compound, drug | Ebselen | ThermoFisher | AAJ63190MA |
| Chemical compound, drug | Erythromycin | ThermoFisher | BP920-25 |
| Chemical compound, drug | Gentamicin sulfate | ThermoFisher | BP918-1 |
| Chemical compound, drug | Kanamycin sulfate | ThermoFisher | BP9065 |
| Chemical compound, drug | Mefloquine | ThermoFisher | AC461130010 |
| Chemical compound, drug | Neomycin sulfate hydrate | ThermoFisher | AAJ6149914 |
| Chemical compound, drug | Tobramycin | Sigma-Aldrich | T4014-500MG |
| Chemical compound, drug | 3-aminobenzoic acid ethyl ester, methanesulfonate salt (tricaine) | Pentair | TRS5 |
| Chemical compound, fluorophore | Texas Red-X succinimidyl ester | ThermoFisher | T20175 |
| Chemical compound, stain | YO-PRO-1 Iodide | ThermoFisher | Y3603 |
| Media | ICS water; Instant Ocean Sea Salt | Spectrum Brands Product SS15-10 | https://www.chewy.com/instant-ocean-sea-salt-aquariums/dp/196228 |
| Media | Middlebrook 7H9 Broth | ThermoFisher | DF0713-17-9 |
| Media | Cation-adjusted Mueller-Hinton Broth (CAMHB) with TES | ThermoFisher | T3462 |
| Hardware | Non-treated 12 and 24 well plates | ThermoFisher | 07-201-589, 07-201-590 |
| Hardware | Non-treated 96 well clear, round bottom plates | GenClone | 25-224 |
| Hardware | Non-treated 96 well black, flat bottom imaging plates | VWR | 37000-550 |
| Hardware | 3D printer filament: PLA | Amazon | B084XS2TS9 |
| Hardware | 125 micron Nylon mesh | ELKO Filtering Co. | 03-125/39 |
| Software | Tinkercad design for netwells |  | https://www.thingiverse.com/thing:5676024 |
| Software | ImageJ | National Institutes of Health | https://imagej.nih.gov/ij/download.html |

### Zebrafish husbandry

All experiments involving live zebrafish were carried out in compliance with Seattle Children’s Research Institute’s (IACUC protocol number ACUC00658) and University of Washington’s Institutional Animal Care and Use Committee guidelines (IACUC protocol number 2997-01). As zebrafish lateral line HCs develop by 5 days post-fertilization (dpf), experiments were conducted when fish were 5 dpf unless otherwise noted. Sex is not determined at this age.

For drug dose response testing, zebrafish of the wildtype genetic background AB and fish that were transgenic for GFP under the *myo6b* HC-specific promoter (Tg(*myo6b:EGFP*)) were raised as described previously^51, 52^. Briefly, zebrafish embryos were collected from 2 hour spawning periods, washed with dilute bleach solution (0.005% sodium hypochlorite) by six hours post-fertilization, and raised in Petri dishes in ICS water (300 mg Instant Ocean/L, 0.56 mM CaCl_2_, 1.2 mM NaHCO_3_)^23^ at a density of 50 larvae per 100-mm^2^ dish, in a dark 28.5 °C incubator until 5 dpf, with dechorionation at 2-4 dpf.

For calcium testing, zebrafish expressing HC-specific GCaMP targeted to the inner mitochondrial matrix were crossed with fish expressing a HC specific calcium indicator to create double-transgenic embryos (Tg (*myo6b:mitoGCaMP3;myo6b:cytoRGECO*)) with both green fluorescent mitochondrial calcium and red fluorescent cytosolic calcium indicators, then raised as described previously^53^. Zebrafish embryos were collected from 2-hour spawning periods and raised in Petri dishes in embryo medium (EM: 14.97 mM NaCl, 500 µM KCl, 42 µM Na_2_HPO_4_, 150 µM KH_2_PO_4_, 1 mM CaCl_2_ dehydrate, 1 mM MgSO_4_, 0.714 mM NaHCO_3_, pH 7.2) at a density of 60 larvae per 100-mm^2^ dish, in a dark 28.5 °C incubator until 5 dpf. Larvae were screened for indicator expression at 4 dpf.

### Drug response testing

Healthy zebrafish lateral line HCs are selectively permeable to fluorescent vital dyes including YO-PRO1, which selectively stains lateral HC nuclei ^13^. The extent of drug-induced injury at different doses can thus be quantified by loss of fluorescent staining in HCs ^13^. Therefore, at 5 dpf, larvae were transferred to 12-well plates (5-11 fish per well, 1 well per condition) containing the relevant compound or combination diluted with ICS water, or ICS water alone or high-dose neomycin to serve as controls. Unless otherwise noted, fish were exposed to drug (either an individual compound or a pairwise combination of compounds) for 4 hours, as a balance between longer exposure times for accommodating varied kill kinetics and shorter exposure times for technical tractability. Treatments with multiple drugs involved co-administering a prepared solution consisting of both drugs with defined concentrations in ICS water. Fish were then washed in fresh ICS water before being transferred to new 12-well plates containing a 2µM solution of YO-PRO-1 for 20 minutes, before being washed in fresh ICS water, anesthetized with tricaine, and transferred to 96-well plates for imaging (1 fish per well, 60 fish per plate due to specifications of the microscope).

To facilitate the transfer of fish between exposures to drugs, fresh ICS, and YO-PRO-1 solutions, we developed custom single-use multi-well baskets (**Figure 1A**), inspired by the baskets described by Thisse and colleagues^54^. We designed the frame for these baskets (https://www.thingiverse.com/thing:5676024), which we printed with a Dremel Digilab 3D45 3-D printer to provide a single point of manipulation for the whole plate. Nylon mesh was melted to this frame as previously described to form a basket for each well of the relevant multi-well plate^54^. Each multi-well basket was disinfected with 10% bleach soak prior to use.

### Imaging and ototoxicity quantification

Fluorescence microscopy was performed using a Keyence BZ-X800 microscope imaging system, which enables high-throughput semi-automated imaging of 96-well plates. Using a GFP filter (525/50nm emission, 470/40nm excitation), each YO-PRO-1-stained fish in the 96-well plate was imaged under a 2x objective to capture the whole organism in brightfield, green fluorescence, and red fluorescence channels.

The resulting images were analyzed for dose-response and drug interaction characteristics with the PEPITA software package, which can be found on GitHub (https://github.com/ma-lab-cgidr/PEPITA-tools). Images did not have to be consistent in exposure or aperture used, as values were postprocessed, making use of the fact that received signal is directly proportional to the exposure time and inversely proportional to the square of the aperture *f*-stop ^55^. Each image was masked using automated object detection to include only the fish itself. Manual masks were created on occasion for avoidance of fluorescent contamination, adjusted segmenting for *myo6b::gfp* fish to remove inner ear HC fluorescence, or exclusion of dead or damaged fish. Each fish was then adjusted for autofluorescence using the fluorophore-free red channel and scored based on the sum of green fluorescence pixel values surrounding 10 of the top 15 brightest pixels in the masked image (**Figure S1**). These scores were standardized by comparison with the median score of wells containing untreated fish and a score representing no remaining HC fluorescence, both derived for each plate. The standardized relative fluorescence unit (RFU) scores were fitted against a log logistic model, as described previously^56^, to estimate dose response properties (e.g. EC_50_, the effective concentration of eliciting 50% of maximal HC damage).

### Aminoglycoside uptake quantification

To further explore the mechanism of macrolide and aminoglycoside antagonism, we fluorescently labeled the aminoglycosides neomycin and gentamicin with Texas Red-X succinimidyl ester to track HC uptake with and without macrolide treatment. To avoid breakdown of fluorescent vital dyes over time, we used larvae with the transgene *Tg(myo6b:EGFP)*; *myo6b::gfp*, that expressed GFP in their HCs. Drug-induced injury was quantified by loss of fluorescence in HCs. Fish were exposed to the fluorescently-labeled drug with or without macrolide co-administration for 30 minutes, before being washed in fresh ICS water and anesthetized with tricaine. Five fish per condition were transferred to 96-well plates for imaging, as per the drug response testing experiments, while another five were fixed in 2% paraformaldehyde, as described previously^16^, to prepare for confocal microscopy.

For the fish transferred to 96-well plates for imaging, fluorescence microscopy was performed using the Keyence BZ-X800. Using the GFP filter and a TexasRed Filter (630/75nm emission, 560/40nm excitation), each *myo6b::gfp* fish was imaged under a 2x objective to capture the whole organism in brightfield, green, and red fluorescence channels. Each fish was then imaged under a 40x objective with z-stacking to track the uptake of the fluorescently labeled aminoglycoside into the HCs in the head of the fish.

To prepare the fish for confocal microscopy, we transferred the fixed fish to fluoromount and mounted them between two glass coverslips. To avoid crushing the fish, a dot of nail polish was placed on each corner of the coverslip so that the top coverslip sat slightly above the bottom coverslip where the fish was mounted. Fluorescent confocal microscopy was performed using a Leica Stellaris microscope. Each fish was imaged under a 40x objective to capture the head of each fish in green and red fluorescence channels. Image analysis was performed in ImageJ.

### FM1-43 uptake quantification

FM1-43 experiments were performed with *myo6b::gfp* fish, raised to 5 dpf as above. Fish were exposed to drug conditions for 30 minutes, with FM1-43 stain added into the drug treatment solution at minute 29, for a one-minute co-treatment stain exposure followed by simultaneous washout of all drugs and stain in fish water and anesthesia in tricaine. Four to seven fish per condition were transferred to 96-well plates for imaging. Using the Keyence BZ-X800 with GFP filter and a custom filter (605/70nm emission, 470/40nm excitation), each fish was imaged at 2x in brightfield, green, and red fluorescence channels as above.

### Lateral line hair cell intracellular calcium quantification

Calcium imaging was performed using an inverted Marianas spinning disk confocal system (Intelligent Imaging Innovations, 3i) with an Evolve 10 MHz EMCCD camera (Photometrics) and a Zeiss C-Apochromat 63x/1.2 numerical aperture water objective. Larvae were 5-6 dpf at the time of imaging. Larvae were first anesthetized in embryo media containing 0.2% tricaine, then stabilized on their sides under a harp, so that posterior neuromasts were exposed to the surrounding media. Imaging was performed at ambient temperature, approximately 25°C. Images were taken every two minutes as z-stacks through neuromasts in 2 µm steps. Neomycin and azithromycin were dissolved as a 4X stock in EM (described in the *<u>Zebrafish Husbandry</u>* section), then added to the bath to achieve final working concentration after a ten-minute baseline. Imaging of neuromasts continued for 50 min. Images were analyzed in ImageJ. Mean fluorescence intensity was measured for individual HCs. For each HC, values were normalized to the average of baseline using Microsoft Excel.

### Bacterial drug response testing

For bacterial drug interaction testing, bacterial strains Staphylococcus aureus ATCC 29213, Escherichia coli ATCC 25922, were grown from frozen stocks on 5% sheep blood agar for 16 hours at 37°C, and Mycobacterium abscessus ATCC 19977 was grown from frozen stocks on Middlebrook 7H10 agar for 3 days at 37°C ^57^ . On the day of inoculation, serial dilutions of the appropriate drugs were performed in separate plastic troughs and then combined in a clear, round bottom, cell-culture treated 96 well plate. Macrolide serial dilutions ascended in concentration row-wise (left to right) and aminoglycoside serial dilutions ascended in concentration column-wise (top to bottom). A bacterial inoculum of 0.5 (±10%) McFarland units was prepared in sterile saline and then diluted 1:100 in the appropriate media. The drug dilution plate was inoculated with 100 µL of the bacterial inoculum, and the plate was sealed and incubated at 37°C for 18 hours to 3 days, depending on the strain. After incubation, plates were read on an indirect mirror box by eye for inhibition of all visible growth and with a SpectraMax i3x microplate reader for absorbance at 600 nm.

### Drug interaction quantification

We quantified drug interactions with the excess over Bliss (EOB) metric ^58^. Given a combination of drugs A and B, EOB was calculated for each individual well exposed to doses of both drugs based on monotherapy and combination responses with the formula *EOB* = *R_a_* × *R_b_* − *R_ab_* (1)^58^. Wells where the expected response (*R_a_* × *R_b_*) and observed response (*R_ab_*) were both less than 10% or greater than 90% were excluded from aggregation to reduce the effect of our choice of dose range. The remaining wells were summarized by averaging into a composite interaction score for the combination; we refer to this metric as the “windowed excess over Bliss” (wEOB).

### Statistical analysis

Error bars, for both point estimates and line plots, indicate 95% confidence intervals, calculated by bootstrapping, as a nonparametric measure of uncertainty ^59^. All zebrafish fluorescence data are included except for dead fish, fish too out of plane to view enough neuromasts for robust analysis, and experiments in which untreated controls are dimmer than several conditions that should have reduced HC fluorescence based on past data or literature. All bacterial data are included except for experiments in which uninhibited bacterial growth controls are less than the 60th percentile of measured wells or media only controls are greater than the 40th percentile of measured wells. Values plotted with no error bars depict individual measurements for the given conditions; when depicted with error bars, points represent the arithmetic mean of all valid measurements. For dose-response curve shift values, effective doses are estimated by interpolation on a logarithmic scale, and confidence intervals are estimated similarly based on bootstrapped point estimates. Shift *p*-values are calculated by extra sum-of-squares F test using the drc package in R ^60, 61^. Elsewhere, *p*-values are calculated with Welch’s unequal variances *t*-test^62^. Multiple testing is adjusted for with Benjamini-Hochberg FDR correction when noted ^63^.

## Code Availability

The PEPITA-tools software package can be found on GitHub at https://github.com/ma-lab-cgidr/PEPITA-tools

## Acknowledgements

We gratefully acknowledge Patricia Wu and Rina Yan for their technical assistance. This work was supported by the National Institute of Health (Grant: R21 DC018341) and the Lura Cook Hull Trust.

## Author contributions

S.M. conceptualized the study and acquired funding. S.M., R.E.H, and D.W.R supervised the project. E.B., E.M., E.M.N, A.M., M.B., A.G., L.G., and N.G. executed the experiments. E.B. developed the analysis pipeline and performed the data analyses. E.B., A.M., H.C.O, D.W.R., R.E.H., and S.M. interpreted the data. E.B., A.M., and S.M. wrote the manuscript and visualized the data. All authors reviewed the draft and assisted in manuscript preparation.

## Notes

### Competing Interest Statement

The authors have declared no competing interest.

