## Supplemental Materials for "*In vivo* screening for toxicity-modulating drug interactions identifies antagonism that protects against ototoxicity in zebrafish"

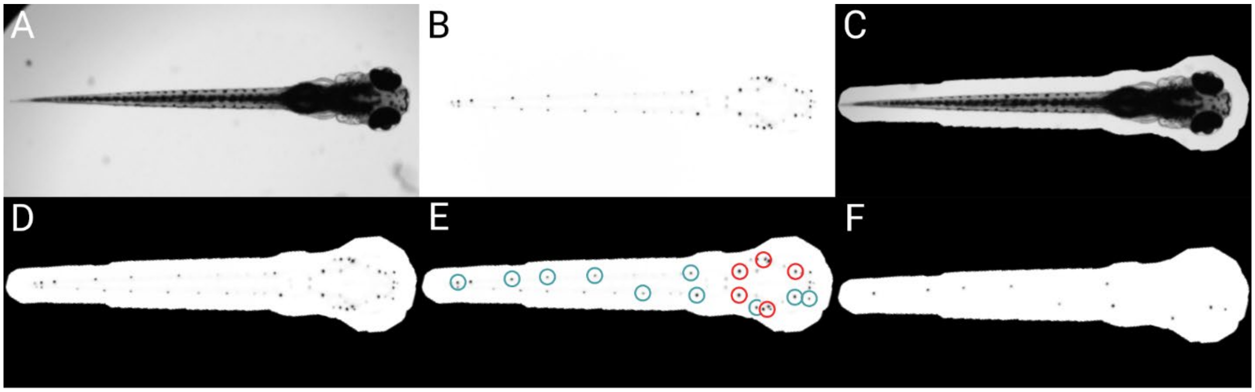


**Figure S1. PEPITA's processing steps for quantifying whole-organism zebrafish image data.** To begin, brightfield (A) and fluorescence (B) images are taken of each organism (here, with neuromasts fluorescently stained with YO-PRO-1). An additional fluorescence channel with no fluorophore present can also be supplied, in which case PEPITA will use it as a baseline to cancel out autofluorescence in the image of fluorescently labeled neuromasts. PEPITA creates a mask (C) by automatically locating the larva by contrast, size, and shape from the brightfield image (which can be overridden manually when necessary). This mask is applied (D) to the fluorescence image in order to identify the points of interest. PEPITA next identifies the 15 brightest local maxima within the masked region (E), excludes the top five (marked here in red), and creates a second mask obscuring everything except small circles (with a radius of 8 pixels by default) around the other ten puncta (marked here in cyan). This second mask is then reapplied to the fluorescence image (F) and the unobscured pixel values that exceed background level are summed to yield the raw fluorescence score for the given larva. Figure created with BioRender.com

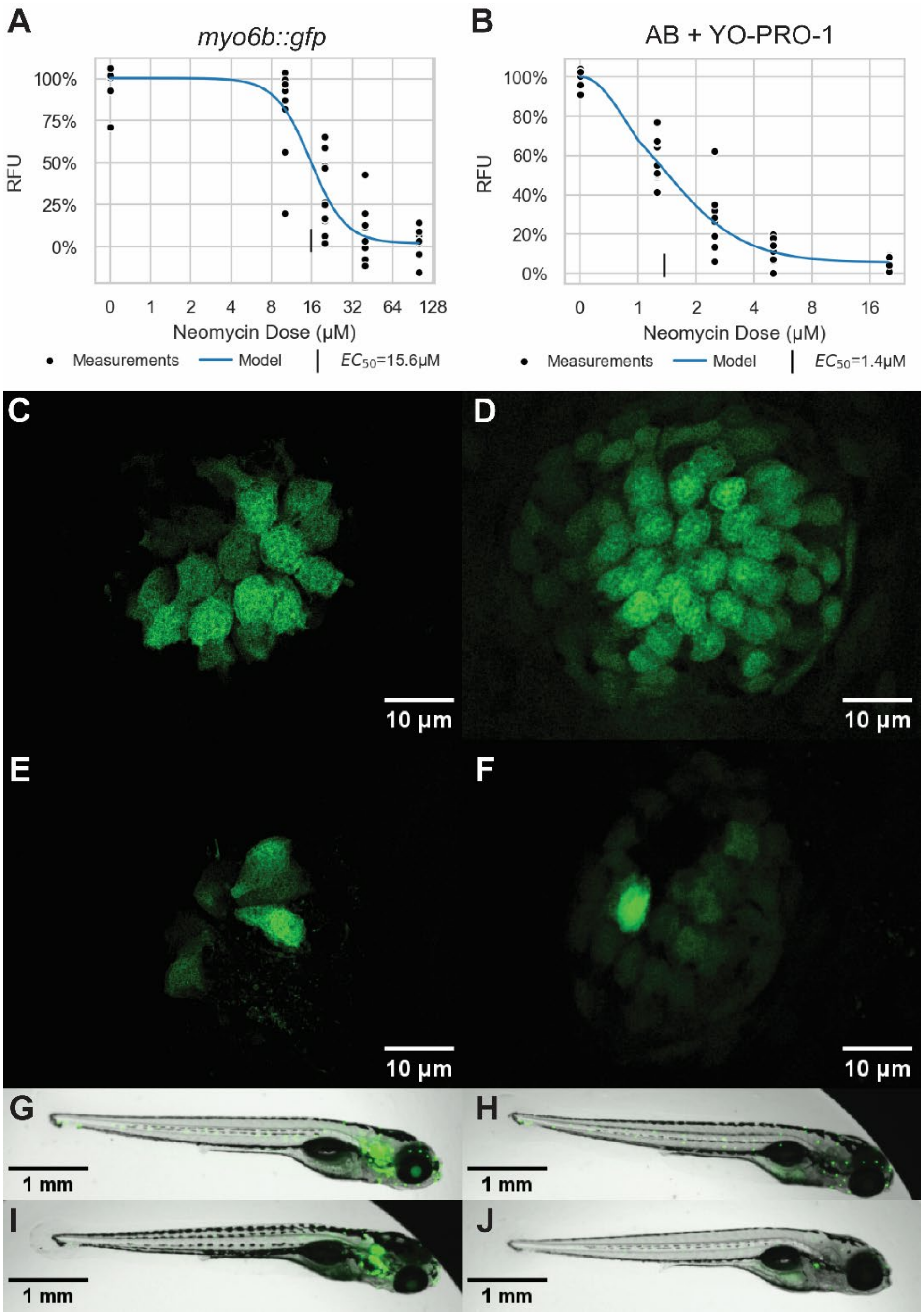


**Figure S2. Comparison between *myo6b::gfp* (left) and YO-PRO-1–stained AB fish (right).** **(A, B)** Representative dose-response curves for each experimental setup when exposed to increasing doses of NEO. **(C, D)** Representative images of undamaged neuromasts in the respective strains. **(E, F)** Representative images of damaged neuromasts in the respective strains, each exposed to about an $EC_{75}$ of NEO (20 μM for *myo6b::gfp*, 2.5 μM for AB). **(G, H, I, J)** Representative images of whole fish, like those PEPITA takes as input. **G** and **H** are untreated; **I** and **J** treated with NEO $EC_{50}$ estimates: 20 μM for *myo6b::gfp*, 2.5 μM for AB. The cluster of fluorescent inner-ear HCs in *myo6b::gfp* larvae requires manual masking for proper quantification with PEPITA.

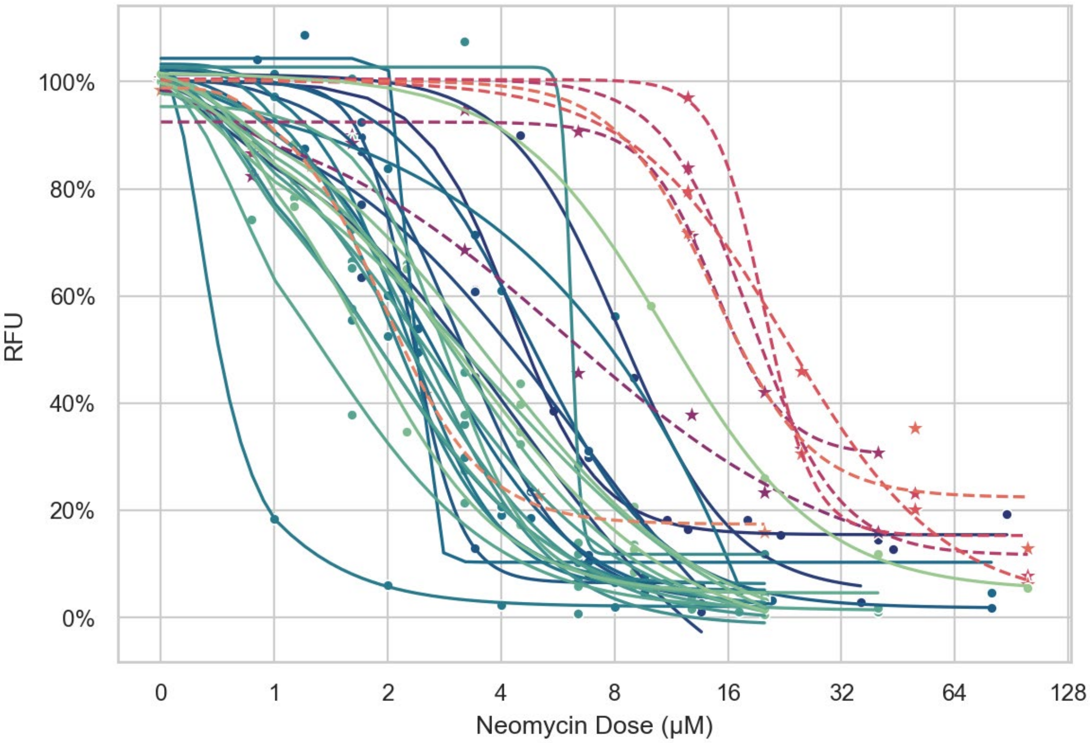


**Figure S3. Overview of the variability in neomycin dose-response curves obtained by PEPITA.** Points are measurements, lines are fitted log-logistic models; circles and solid lines represent AB fish stained with YO-PRO-1, stars and dashed lines represent *myo6b::gfp* fish.

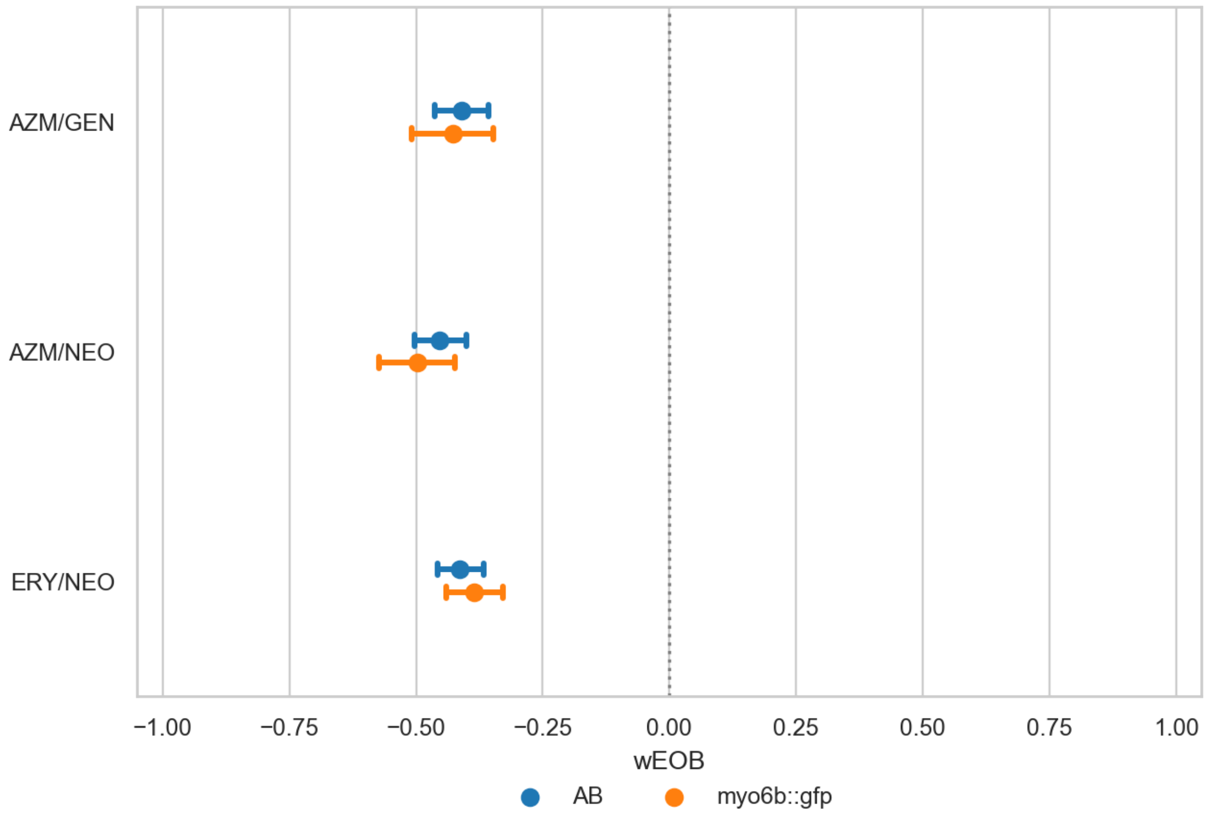


**Figure S4. Comparison of aggregate wEOB value between strains.** The aggregate wEOB for each combination tested in both strains shows strong antagonism and does not significantly differ between strains (AZM/GEN, p=0.75; AZM/NEO, p=0.35; ERY/NEO, p=0.44).

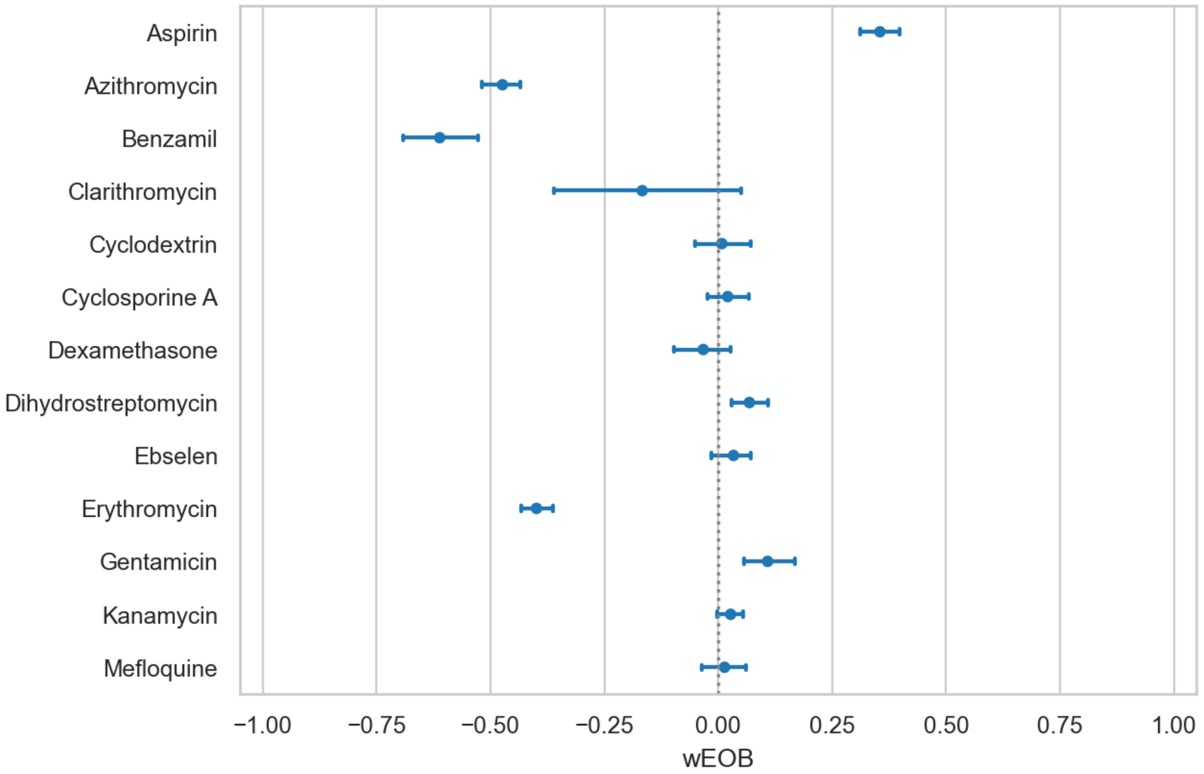


**Figure S5. Overview of interaction scores observed between neomycin and various other drugs screened for significant interactions.** Increasingly negative wEOB scores indicate increasing antagonism between NEO and the listed drug; increasingly positive wEOB scores indicate increasing synergy. Most drugs lie near zero, as would be expected for noninteractive compounds. The only listed compounds that deviate toward antagonism are the three macrolides tested, plus benzamil, a potent MET channel inhibitor well established in antagonizing NEO uptake.

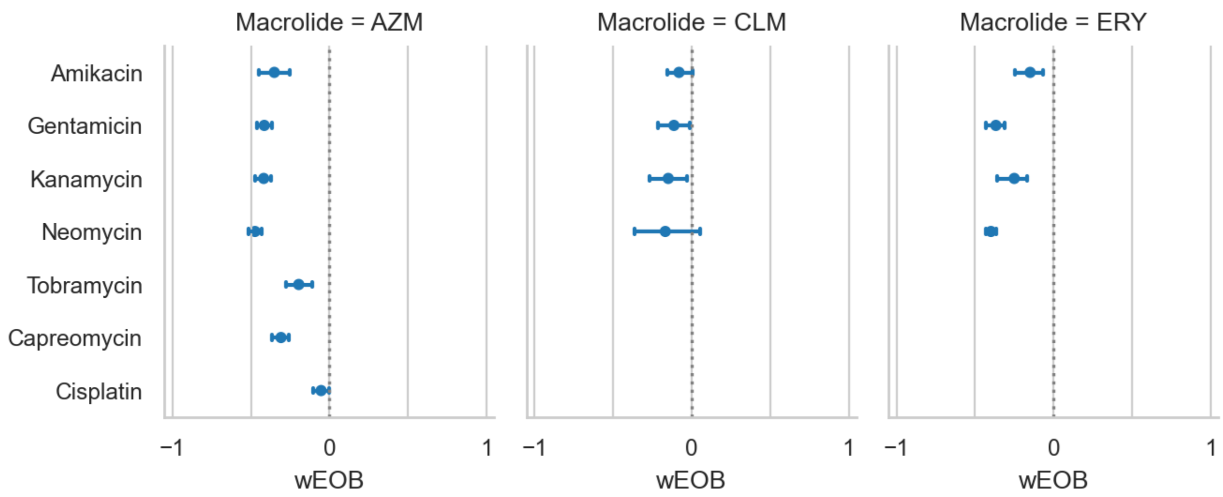


**Figure S6. Overview of interaction scores between macrolide antibiotics and ototoxic drugs tested.**

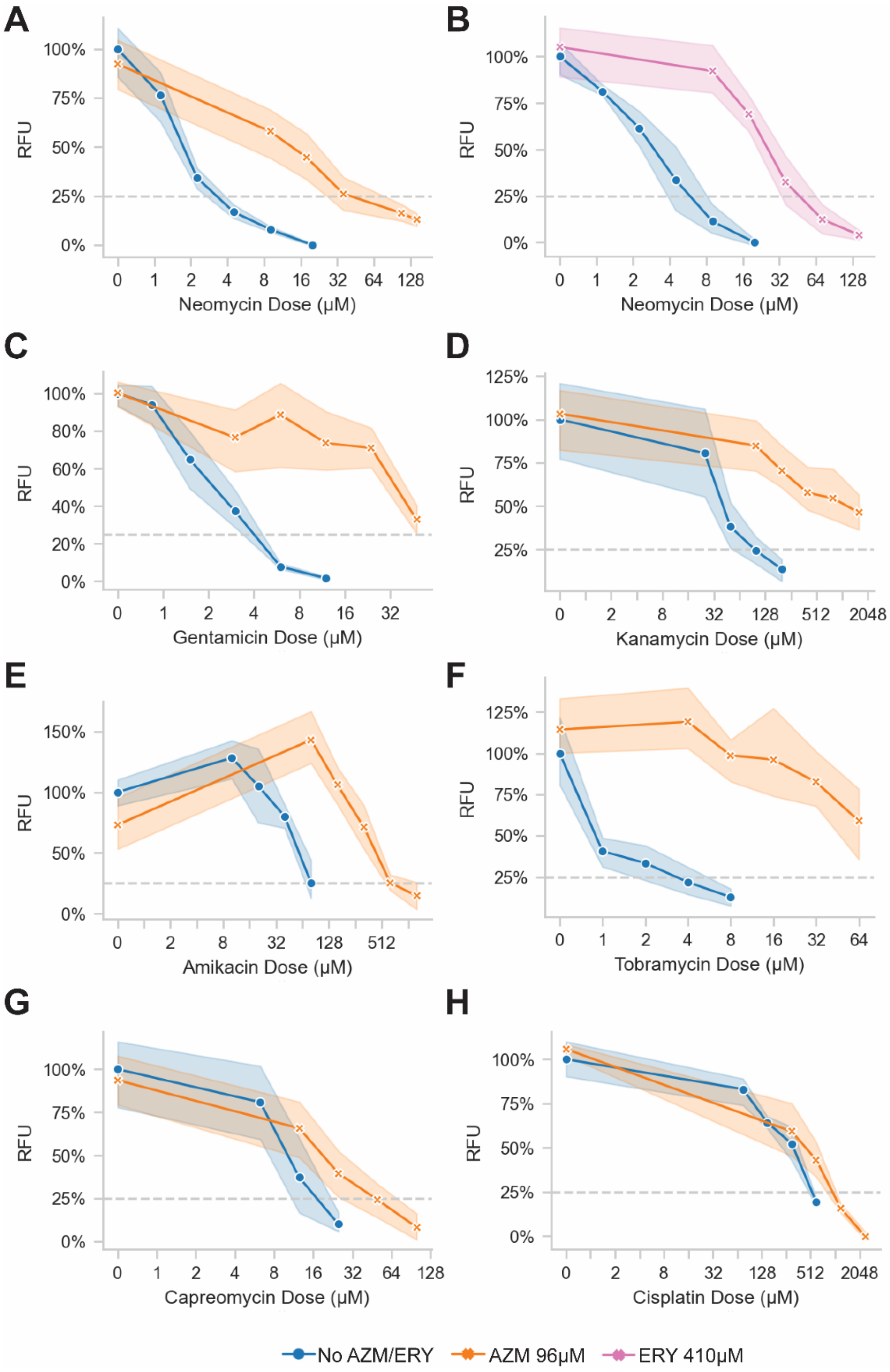


**Figure S7. AZM antagonizes aminoglycoside-induced ototoxicity.** We observe at least 8-fold increase in $EC_{75}$ across all compound combinations tested. Non-aminoglycoside ototoxins CAP and CIS are antagonized to a lesser extent. ERY antagonizes NEO-induced ototoxicity to a comparable extent to AZM.

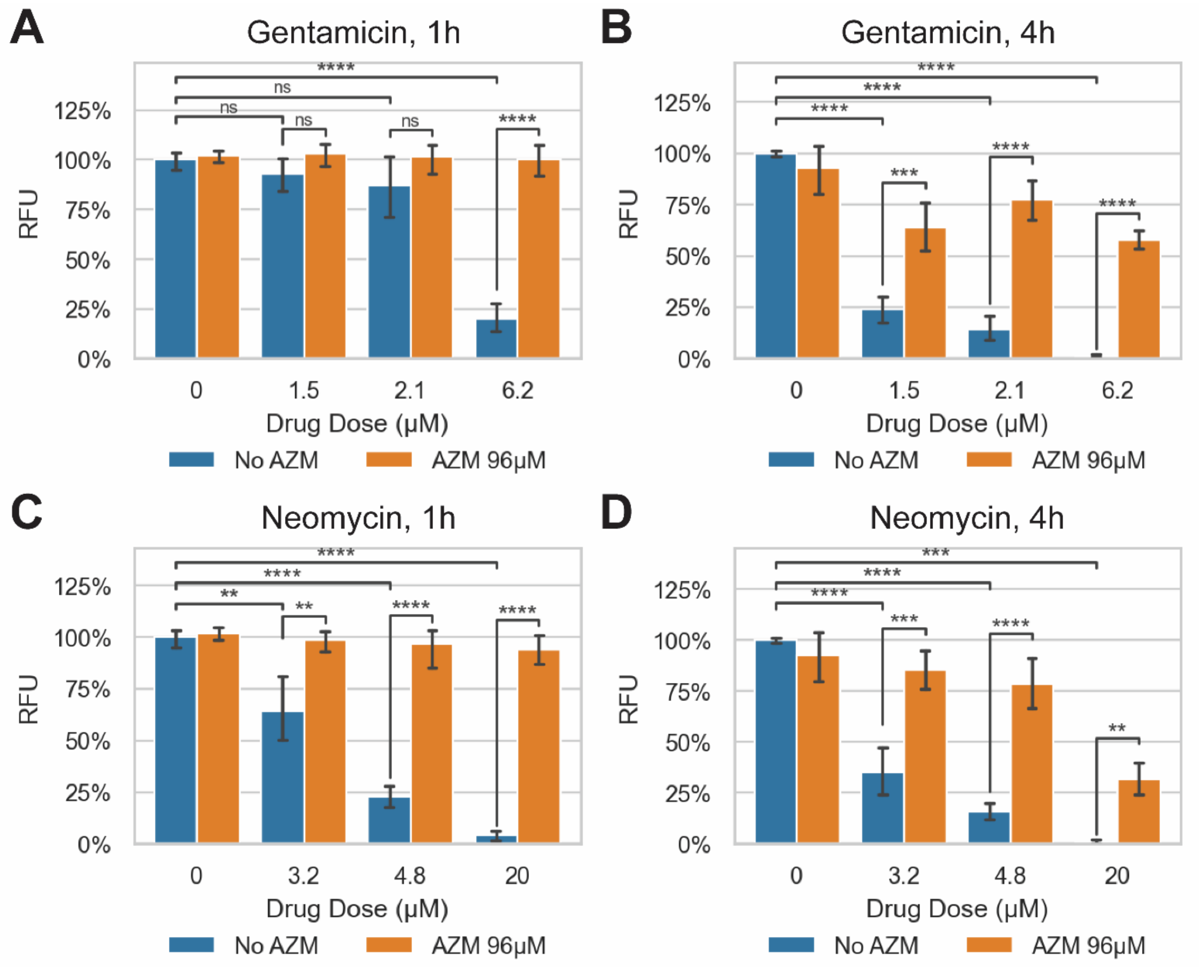


**Figure S8. Extent of aminoglycoside-induced ototoxic damage and macrolide-conferred otoprotection as a function of time.** Doses were chosen to represent estimated effective concentrations necessary to achieve 50%, 75% and 99% inhibition, respectively, as calculated from previous experiments at the 4-hour timepoint. AZM confers significant protection in all conditions where monotherapy shows significant damage as compared to untreated control. Corrected p-values: * = $p < 0.05$; ** = $p < 0.01$; *** = $p < 0.001$; **** = $p < 0.0001$.

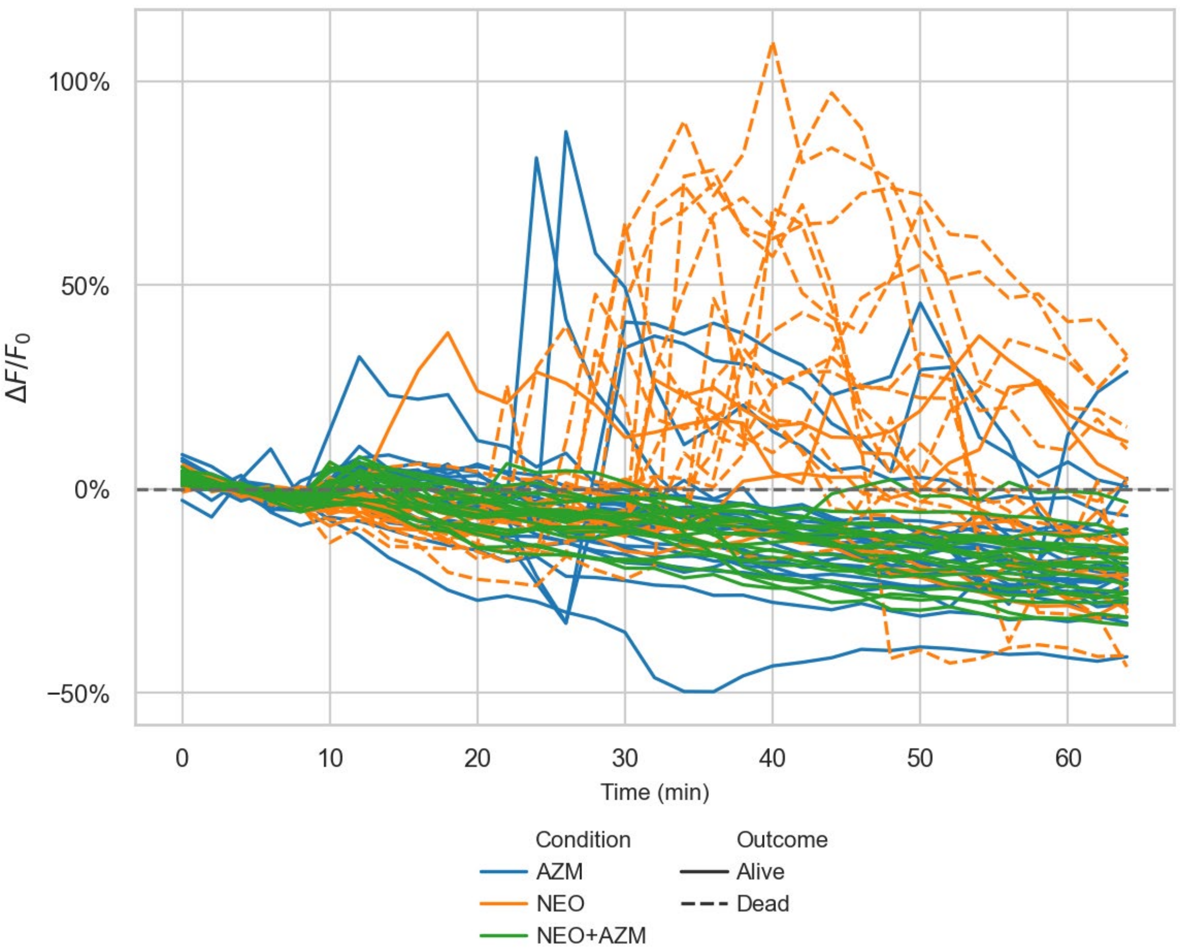


**Figure S9. Quantification of mitochondrial $Ca^{2+}$ levels in individual hair cells in response to AZM treatment (blue), NEO treatment (orange), or the two co-administered (green), as measured by mitoGCaMP fluorescence signal**. Drugs were administered at t = 10 minutes. A majority of cells exposed to NEO alone went on to die (14/21 cells from 9 neuromasts in 4 fish), while most exposed to AZM alone (24/25 cells from 11 neuromasts in 5 fish) and all treated with both drugs (19/19 cells from 10 neuromasts in 5 fish) survived.

| ***Uptake Inhibitor*** | ***Name*** | ***Details*** |
|---|---|---|
| 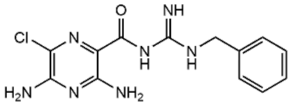 | *Benzamil* | *MET channel inhibitor (PMID 22967486)* |
| 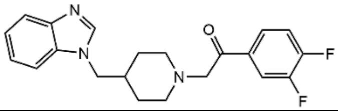 | *UoS-7692* | *MET channel inhibitor (PMID 33735112)* |
| 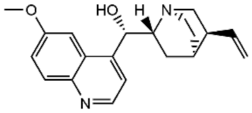 | *Quinine* | *MET channel inhibitor (PMID 15181168)* |
| 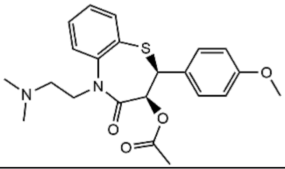 | *Diltiazem* | *MET channel inhibitor (PMID 15181168)* |
| 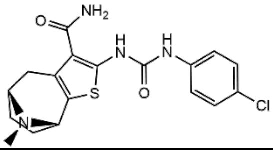 | *ORC-13661* | *MET channel inhibitor (PMID 31391343)* |
| 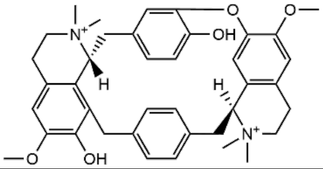 | *Curare* | *MET channel inhibitor (PMID 15181168)* |
| 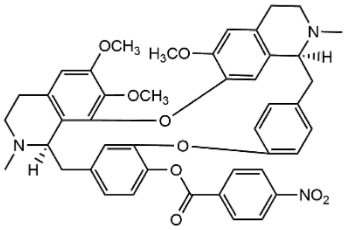 | *E6 Berbamine* | *MET channel inhibitor (PMID 27065807)* |
| 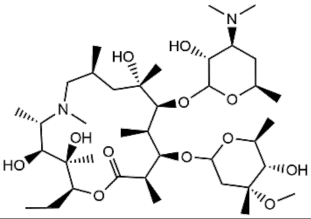 | *Azithromycin* | *Prevents neomycin uptake by means other than MET channel inhibition (this study)* |
| 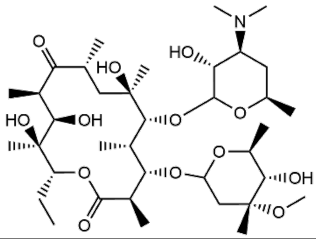 | *Erythromycin* | *Prevents neomycin uptake by means other than MET channel inhibition (this study)* |

**Table S1. Comparison between macrolides and a range of known MET channel blockers.**
